# Investigating low-velocity fluid flow in tumours using convection-MRI

**DOI:** 10.1101/200279

**Authors:** Simon Walker-Samuel, Thomas A. Roberts, Rajiv Ramasawmy, Jake Burrell, S. Peter Johnson, Bernard Siow, Simon Richardson, Miguel Gonçalves, Douglas Pendsé, Simon P. Robinson, R. Barbara Pedley, Mark F. Lythgoe

**Author notes:** Corresponding author: Simon Walker-Samuel, Address: Centre for Advanced Biomedical Imaging, University College London, Lower Ground Floor, Paul O’Gorman Building, 72 Huntley Street, London, WC1 6DD. The authors declare no potential conflicts of interest.

## Abstract

Several distinct fluid flow phenemena occur in solid tumours, including intravascular blood flow and interstitial convection. To probe low-velocity flow in tumors resulting from raised interstitial fluid pressure, we have developed a novel magnetic resonance imaging (MRI) technique named convection-MRI. It uses a phase-contrast acquisition with a dual-inversion vascular nulling preparation to separate intra- and extra-vascular flow. Here, we report the results of experiments in flow phantoms, numerical simulations and tumor xenograft models to investigate the technical feasibility of convection-MRI. We report a good correlation between estimates of effective fluid pressure from convection-MRI with gold-standard, invasive measurements of interstitial fluid pressure in mouse models of human colorectal carcinoma and show that convection-MRI can provide insights into the growth and response to vascular-targeting therapy in colorectal cancers.

## Introduction

The convection of fluid in biological tissue is normally associated with intravascular blood flow, in which fluid flows from high to lower pressure, at velocities ranging from 5 – 70 cm s^−1^ in arteries (1,2), 0.8-2.5 mm s^−1^ in capillaries (3) and 1.5 – 28 cm s^−1^ in veins (1,2). Fluid within the interstitium is normally maintained at or just below atmospheric pressure by venous and lymphatic vessels (4), and several pathological conditions, including cancer, can disrupt key pressure regulation processes, leading to widespread elevated interstitial fluid pressure (IFP) (5).

Raised IFP in solid tumors has been the subject of extensive research to investigate its pathophysiological consequences, which can include an increased metastatic potential (6–8), higher rates of proliferation (9,10) and impaired drug delivery (11–16). It is associated with numerous pathophysiological processes in tumors, and has been described as a ‘universal generic early-response marker of tumor response to therapy’ (17). A physical consequence of raised IFP in tumors is the slow convection of fluid through the interstitium (4), which has been widely reported and studied (4,18–21), but is technically challenging to measure, and has only been directly assessed through invasive means (22). IFP is normally elevated in the center of tumors and maintained close to atmospheric pressure at the periphery by surrounding tissues, resulting in the convection of fluid from the center outwards (21), with a velocity in the range 0.1 to 50 μm s^−1^ (18).

In this study, we report the development of a new, non-invasive imaging technique that aims to probe the slower-flowing components of fluid flow in tumors; in particular, those associated with interstitial fluid. It is based on velocity-encoded MRI, with a dual inversion preparation that aims to null the faster-flowing vascular signal. We describe our approach in this study, which we have named convection-MRI, assess its technical feasibility and evaluate its underlying assumptions. Specifically, convection-MRI assumes that: 1) the intravascular signal can be completely nulled by dual inversion pulses; 2) velocity encoding is sufficiently sensitive to measure the velocity of fluid flow in the interstitium; and 3) the influence of nulled fluid exchanging between intra-and extra-vascular compartments is negligible. The physical principles underlying the acquisition are described schematically in **Supplemental Movie 1** (https://goo.gl/uQ5oZp). Both the dual inversion and velocity-encoding techniques have been used separately in other contexts (for example, references (23–25) for the dual inversion preparation and references (26,27) for velocity-encoding), but their combined use is novel in the current application, and neither has been used to study interstitial flow.

The study was undertaken in two distinct mouse xenograft models of colorectal cancer, both during their growth and following treatment with a vascular disrupting agent (VDA). The results of this multiparametric, noninvasive MRI analysis allowed numerous novel insights into the differential growth characteristics of two colorectal cell lines with differing vascular and cellular morphology (SW1222 and LS174T) to be gleaned, and how these relationships were modified with VDA therapy. SW1222 cells form well-differentiated tumors with glandular structures resembling normal colon, and are well-vascularized (28), with minimal hypoxia (29,30). In contrast, LS174T tumors are poorly differentiated, with heterogeneously distributed vasculature (including avascular regions) (28), and regional hypoxia (29,30), and so provide a useful comparative model system.

## Materials and Methods

### Convection-MRI

A sequence diagram for the convection-MRI sequence is shown in **Supplemental Figure 1**, which was implemented on a 9.4 T horizontal bore MRI scanner (Agilent, Santa Clara, California), with a maximum gradient strength of 1 T m^−1^. The sequence consisted of two modules: a dual inversion-recovery preparation that aimed to null the signal from flowing blood, followed by velocity-encoding. The first adiabatic inversion pulse in the vascular nulling preparation was applied globally and was followed immediately by a slice-selective inversion (thickness 0.3 cm). Inversion pulses were 2 ms in duration, and were separated by 4 μs. A *T*_1,blood_ of 1900 ms was assumed, which was taken from previous measurement in the mouse ventricular pool (31). At a time *t*_*rec*_ following the global inversion pulse, the signal from fluid in blood vessels passes a null point, where *t*_*rec*_ = ln(2)*T*_1,blood_ (giving *t*_*rec*_ = 1317 ms for the value of *T*_1,blood_ used here).

Following *t*_*rec*_, a standard single-slice gradient echo readout sequence was applied (TE, 2.6 ms; TR, 2500 ms; flip angle, 30°; slice thickness, 1 mm; field of view, 35×35 mm2, matrix size, 128×128). The v_*enc*_ parameter is the maximum velocity that can be measured without producing phase aliasing, and depends on the amplitude and duration of velocity encoding gradients (32). In this study we used gradient amplitude (G) of ±300 mT m^−1^ and gradient duration (*τ*) of 3 ms to give a v_enc_ of 2200 μm s^−1^. Velocity-encoding required two repetitions of the sequence, the second of which used bipolar gradients of opposite polarity to the first. The difference in the phase shift induced by the gradients, between the two measurements, Δ*ϕ*, is proportional to fluid velocity, according to the following relationship:

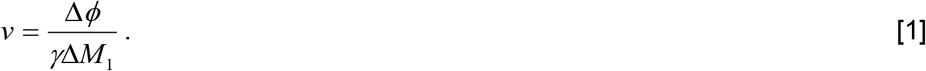

Here, *v* is fluid velocity, γ is the proton gyromagnetic ratio (267.513 × 10^6^ rad s^−1^ T^−1^) and Δ*M*_1_ is the difference in first order velocity-encoding gradient moments (where v_*enc*_ = π/γΔ*M*_1_ and Δ*M*_1_ = 2G*τ*^2^).

The convection-MRI sequence was used to measure fluid velocities in two directions, corresponding to phase and readout imaging gradient orientations. Total scan duration was 25 minutes. In-plane fluid velocity vectors were constructed for each pixel and displayed using in-house software written in Interactive Data Language (ITT, Boulder, Colorado). A streamlining algorithm (iVector) was used to visualise pathways between neighbouring velocity vectors, which used points with low velocity relative to the distribution within the image as seed points.

### Mouse tumor models and in vivo MRI

All *in vivo* experiments were performed in accordance with the local ethical review panel, the UK Home Office Animals Scientific Procedures Act (1986) and UK National Cancer Research Institute (NCRI) guidelines (33). Six-week old MF1 *nu/nu* mice were injected subcutaneously on the midline of the right flank, close to the hind limb (to minimize respiration motion during imaging), with 5×10^6^ SW1222 or LS174T cells.

MRI was performed following 10 to 15 days’ growth (giving a tumor volume of approximately 0.6 cm^3^). Immediately prior to scanning, anesthesia was induced in the mice with 3-4% isoflurane carried in O_2_. Once anaesthesia was confirmed, mice were transferred to a bespoke imaging cradle, in which anesthesia was maintained with 1.25 to 1.75% isoflurane, which was delivered via a nose cone. Mice were positioned on their side, with tumor uppermost. Tumors were liberally covered in dental paste (Charmflex, Dentkist, South Korea), to eliminate bulk motion. When set (typically taking 2-3 minutes), the dental paste held the subcutaneous tumor tissue in place via contact with the imaging cradle. Once prepared, the cradle was positioned in the center of the 9.4T MRI scanner. Throughout imaging, core body temperature was monitored and maintained at 36.5 °C using a warm air blower and physiological monitoring apparaturs (SA Instruments, Stony Brook, NY). Respiratory rate was also monitored and maintained at 40 to 60 respirations per minute by varying the isoflurane fraction. RF pulses were transmitted and received with a 39 mm birdcage coil (Rapid MR International, Columbus, OH). At the end of *in vivo* experiments, mice were culled via cervical dislocation, and tumor tissue was harvested for further analysis.

### Determining the sensitivity of convection-MRI to low-velocity flow

To evaluate the ability of the 9.4 T MRI scanner to measure the velocity of fluid travelling at velocities of the order of 1 – 100 μm s^−1^, an imaging phantom was constructed from a 5 mL syringe, with an internal diameter of 11 mm (see **Figure 1a**). This simple phantom provided easily controllable fluid velocities, with coherent, laminar flow profiles, that can be compared against convection-MRI velocity measurements, to inform subsequent *in vivo* experiments. A syringe driver was used to inject water into the phantom, via silicon tubing that incorporated a 3 m loop within the scanner bore to allow the flowing water to become fully magnetised. Convection-MRI data were acquired with no vascular nulling (i.e. equivalent to a purely velocity-encoding sequence) from a sagittal slice through the chamber of the phantom. The inflow rate of the phantom was varied between 0.007 and 6.85 mL min^−1^ (giving a theoretical fluid velocity range of 1 to 1000 μm s^−1^, assuming laminar flow). Convection-MRI data were acquired in a cross-section across the phantom, without the vascular nulling preparation. *v_enc_* for these acquisitions was varied between 1000 and 5000 μm s^−1^, applied in separate acquisitions in the read and phase encoding directions.

**Figure 1.**
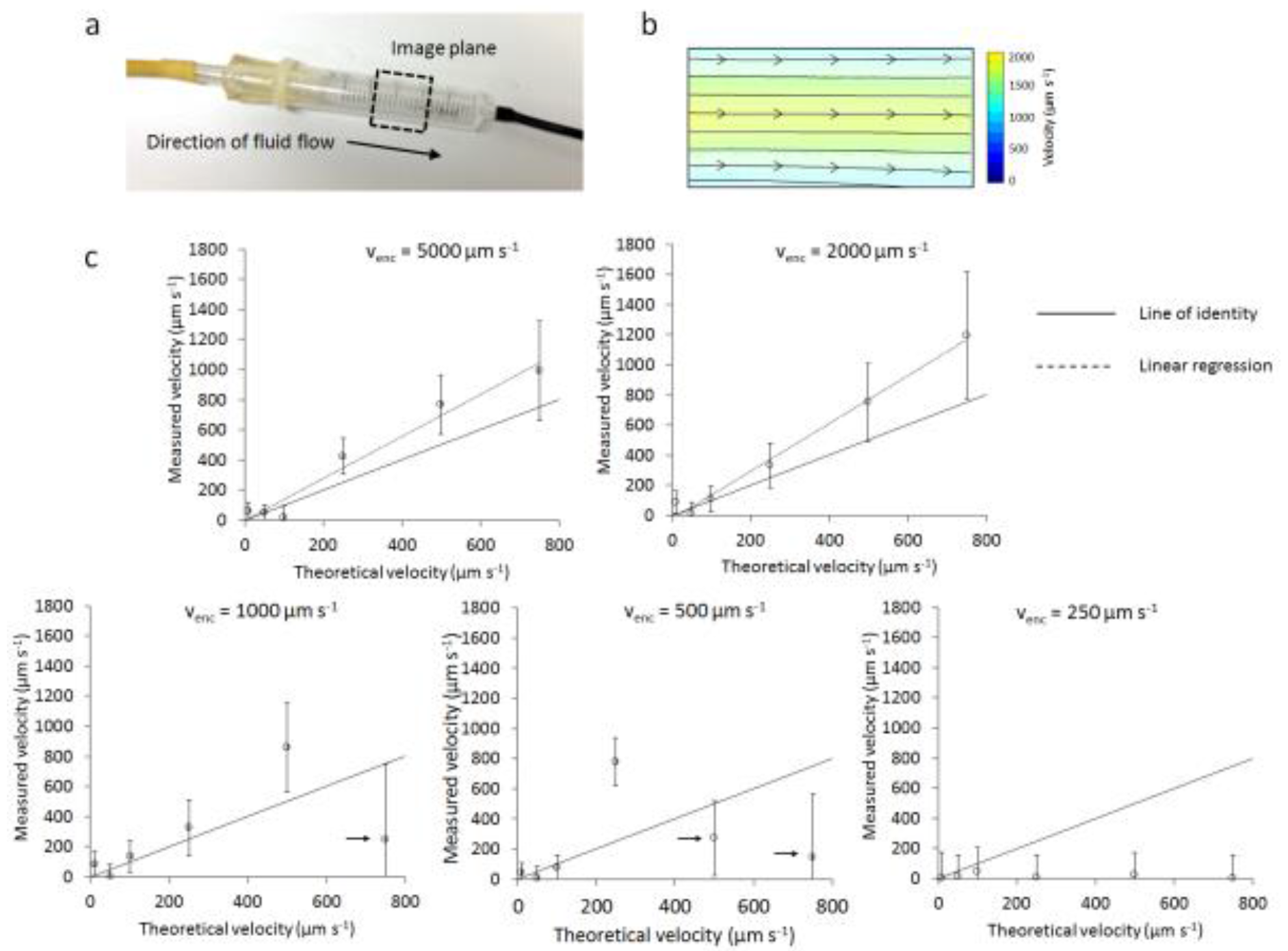
Evaluation of fluid velocity measurements in a flow phantom. (a) Photograph of the fluid velocity flow phantom, based on a 5 mL syringe. The black, dashed box shows the approximate location of the imaging plane used. (b) Fluid velocity vector field, acquired from a slice through the flow phantom, color-coded to reflect the fluid speed, and with the direction characterised using a streamlining algorithm (black lines). (c) The average velocity in the center of the phantom is shown plotted against the inflow rate (error bars show the standard deviation in each measurement), for *v_enc_* ranging from 5000 to 250 μm s^−1^. A significant linear correlation was measured for *v_enc_* values of 5000 and 2000 μm s^−1^ (*p* < 0.01). At small *v_enc_*, aliasing was noted at higher inflow rates (marked with arrows), alongside signal crushing (particularly evident at *v_enc_ =*250 μm s^−1^). Each data point on the graph corresponds to the average value measured in 20 to 30 voxels in the phantom.

### Evaluation of the efficacy of vascular nulling

Phantom and *in vivo* experiments were performed to evaluate the ability of the convection-MRI sequence to null the signal from fluid flowing within tumor blood vessels. Two sets of images were acquired: the first without the vascular nulling (dual inversion) preparation, and the second with vascular nulling. Taking the ratio of images acquired with and without vascular nulling, referred to as the nulling ratio (*n_v_*), enabled vascular volume to be approximately estimated (23).

### Influence of the assumed value of blood T_1_ on vascular nulling in tumors

In a separate cohort of mice bearing SW1222 or LS174T tumors (n = 3 for each), the effect of varying *t_rec_* (equivalent to varying the assumed value of *T*_1,blood_) was investigated. Nulling ratio maps were produced in each tumor for 17 values of *t_rec_*, ranging from 100 to 5000 ms. The average and standard deviation of the nulling ratio was estimated across the cohort and plotted as a function of *t*_*rec*_.

### Estimation of apparent fluid pressure (P_eff_) from convection-MRI data

IFP was estimated from velocity vector fields using Darcy’s law:

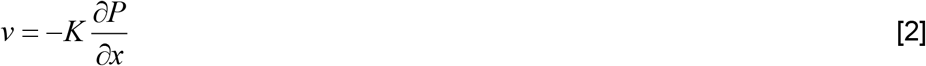

where *K* is the hydrostatic conductivity, *v* fluid velocity and *P* fluid pressure. To estimate *P*, velocity vector fields, measured using convection-MRI, were numerically integrated using a finite difference approach. To define absolute pressure, rather than relative changes in pressure, estimates were offset by the average value at the periphery of the tumor, with the assumption that the periphery of the tumor is in equilibrium with surrounding normal tissues (34). We refer to this as the effective fluid pressure (*P_eff_*). Thus, a boundary condition for the model was that the mean pressure at the periphery of the tumor is zero. For comparison, direct measurement of IFP was performed using a clinical pressure transducer (Stryker, Kalamazoo, Michigan, USA), modified to incorporate a 28G needle (34). Samples were acquired at 8 to 12 locations within each tumor, at depths corresponding to the convection-MRI imaging plane.

### The influence of fluid exchange and incomplete vascular nulling on interstitial fluid velocity measurements

To further evaluate nulling efficiency and the influence of nulled fluid entering the interstitium on convection-MRI IFV estimates, a series of simulations were performed in the Interactive Data Language (IDL, Excelis, Boulder, Colorado). A multi-compartment model of a vascular tumor was constructed with a blood volume (*f_v_*) of 8%, intracellular volume (*f_ic_*) 80% and the volume of the extracellular extravascular space (EES, *f_ees_*) 12%, reflecting those values previously measured in SW1222 colorectal carcinoma mouse xenograft tumors (35). A single voxel within a section of tumor was modelled, with a slice thickness corresponding to that used in the convection-MRI sequence.

Tumor blood vessels were modelled as randomly oriented cylinders. The radius of each vessel, *r*, was generated by randomly drawing from a distribution of vessel radii previously reported by Burrell *et al* (35), which we fitted to an exponential probability distribution (p(*r*) = 0.15 exp(−70*r*)). The total blood vessel volume in each simulation was constrained to be within 10% of the average value (i.e. *f_v_* ranged from 7.2 to 8.8%). The pressure drop, Δ*P*, across each vessel was fixed at 2 mm Hg mm^−1^ and blood viscosity, *η*, was fixed at 3.5×10^−3^ N s m^−2^. Blood flow, *F*, in each vessel was calculated according to the Hagen-Poiseuille equation:

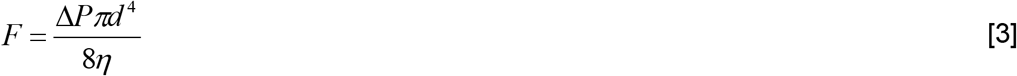

from which blood velocity, *V*_v_, was given by *V*_v_ = *F* / *d*^2^, where *d* is the vessel diameter. The change in signal phase induced by fluid flowing in each blood vessel, in the presence of velocity-encoding gradients in the *x*, *y* or *z* directions (and with zero baseline phase), was given by

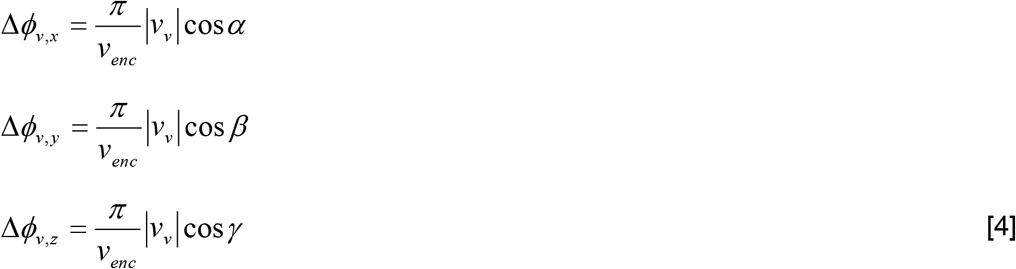

where |*V_v_*| the magnitude of the blood velocity in the vessel, and *α, β* and *γ* are the angles of the vessel, relative to the *x*, *y* and *z* imaging gradient axes, respectively. The total signal from all blood vessels, for velocity-encoding gradients applied in the *x*-direction (the only direction evaluated in the simulations, for simplicity), was then

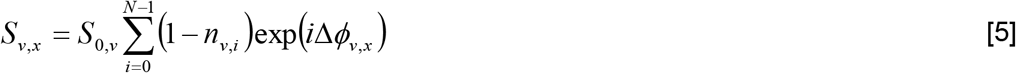

*S*_0,v_ was fixed at unity and relaxation effects were not modelled. To model the effect of fluid nulling in the vascular compartment a factor *n*_v_ was introduced, which ranged from 0 and 1, representing zero to complete nulling. *n*_v_ was calculated by the fractional volume of nulled fluid that replaced un-nulled fluid in blood vessels, at the end of the recovery time following the dual inversion pulses. This was given by

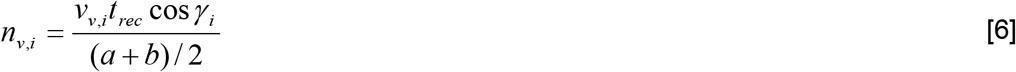

where *a* is the thickness of the imaging slice (1 mm) and *b* is the thickness of the slice-selective inversion pulse (3 mm).

Interstitial fluid velocity was varied between 1 and 100 μm s^−1^ and was directed along the x-axis, for simplicity. By analogy with the vascular signal, the signal from the interstitial compartment, for velocity-encoding gradients applied in the *x* direction, was

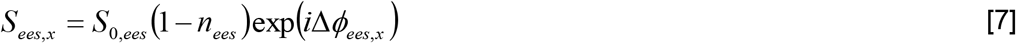

where 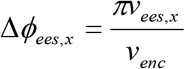 and *S*_0,ees_ was fixed at unity. Simulated nulling of EES fluid was performed explicitly, with *n*_ees_ ranging from 0 to 1. The total MR signal *S* was then given by

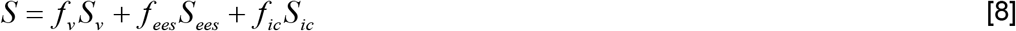

with *f_v_* + *f_ees_* + *f_ic_* = 1 and *S*_ic_ = 1.

1000 Monte Carlo iterations were performed, within each of which a different distribution of blood vessels (drawn from the parent distribution described above) was simulated. As a final step, Gaussian noise was added to each set of simulated data to give signal-to-noise characteristics comparable to those found in *in vivo* data. The phase of data simulated with velocity-encoding gradients applied in the *x*-direction was then estimated, converted into a velocity estimate, and compared with the simulated EES velocity.

Two sets of simulations were performed. Firstly, the effect of nulled fluid entering the interstitium was investigated by varying *n_ees_* between 0 (no nulling) and 1 (complete nulling). Intravascular nulling, *n_v_*, was calculated according to Eq. 5, allowing the theoretical efficacy of vascular nulling to also be determined. In the second set of simulations, to ascertain the influence of un-nulled fluid in the vascular compartment on convection-MRI measurements, *n_v_* was explicitly scaled between 0 and 1, and *n*_ees_ was fixed at 0.

### Arterial spin labelling

Arterial spin labeling (ASL) data were acquired using a flow-sensitive alternating inversion recovery (FAIR) Look-Locker ASL sequence, with a single-slice spoiled gradient echo readout (36,37) (echo time, 1.18 ms; inversion time spacing, 110 ms; first inversion time, 2.3 ms; 50 inversion recovery readouts; 4 averages, all geometric parameters matched to the convection-MRI acquisition). FAIR-ASL does not require a labelling artery to be identified, and instead uses two acquisitions with different inversion preparations: the first, a global inversion pulse, inverts all spins; the second a localized, slice-selective pulse inverts spins within the tissue of interest (a 3 mm slice thickness was used here). The difference between these two signals can be used to measure perfusion, and here regional perfusion maps were calculated as described by Belle *et al.* (38), with an assumed blood-partition constant of 0.9. As for convection-MRI measurements, the *T*_1_ longitudinal relaxation time of blood was assumed to be 1900 ms (39). FAIR-ASL data were acquired in a single slice located in the same position as convection-MRI data.

### Diffusion MRI

Diffusion MRI data were acquired using a fast spin-echo sequence with geometry matched to convection-MRI data. The sequence contained the following parameters: repetition time (TR), 1500 ms; echo train, 4; echo time spacing, 7.74 ms; 5 b-values (150, 300, 500, 750, 1050 s mm^−2^, all applied in the slice-select direction). Diffusion-weighted (magnitude) data were fitted to a single exponential model of the form

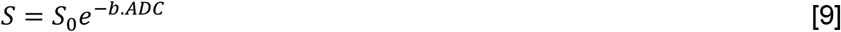

on a pixel-by-pixel basis, using a Bayesian maximum a posteriori approach that takes into account the Rician distribution of magnitude MRI data (40). *b* is the b-value associated with each diffusion-weighted acquisition, and *S* is the acquired signal magnitude for each pixel. Maps of each fitted parameter (the apparent diffusion coefficient (ADC) and *S*_0_) were generated.

### Dynamic contrast-enhanced MRI

Gadolinium-DTPA (Magnevist, Bayer, Leverkusen, Germany) was prepared to a concentration of 0.2 mM (factor of 10 dilution) and transferred to a 50 mL Falcon tube. This was placed in a power injector (Harvard Instruments, Cambourne, UK), positioned inside the scanner room, with a 2 m tube attached, feeding into an 18 G plastic cannula. The dead volume of the tubing was primed with contrast agent and the cannula positioned within the intraperitoneal cavity. Contrast agent was infused at 0.005 mmol kg^−1^ min^−1^ for the duration of the scan (total dose 0.2 mMol kg^−1^). A gradient-echo sequence was used to acquire three-dimensional contrast-enhancement data (TE, 2.43 ms; TR, 107 ms; flip angle 20°; 20 slices; slice thickness 0.5 mm; matrix size, 128×128; FOV, 35×35 mm; temporal resolution 12.8 s; 2 averages), both prior to the start of the infusion, and 40 minutes later. The change in signal intensity induced by contrast agent was calculated by subtracting the baseline signal from the post-infusion data, on a pixel-by-pixel basis, and expressed as a percentage. Contrast enhancement under this infusion protocol will depend on the volume accessible to Gd-DTPA and interstitial transport characteristics (e.g. interstitial fluid pressure and conductivity) (34).

### Microvascular casting

Microvascular casting was performed for colocation of the convection-MRI and perfusion MRI with the tumor vasculature. Casting was performed immediately following MRI. Briefly, the heart was exposed and a blunted 25G butterfly cannula was inserted through the heart and into the ascending aorta. 10 mL of saline was injected through the cannula using a syringe driver (Harvard Instruments, MA, USA). Once the saline injection was complete, casting material (Microfil, Flowtech, Boston, USA) was injected via the same cannula at 60 mL hr^−1^. The microfil used was colored yellow and could be seen entering the vascular bed of the subcutaneous tumors; perfusion of resin was continued until no further coloration of tumor vasculature occurred (typically around 25 mL of resin in total, of which a significant proportion is lost as waste). Following perfusion, the resin was allowed to completely polymerise, which took approximately two hours. Tumors were then carefully resected from the mouse carcass, mounted in chlorine-free plastic food wrap inside a 20 mL syringe (Sigma-Aldrich, Gillingham, Dorset), with plunger removed, and sent for micro-CT.

Micro-CT was performed on a Skyscan 1172 (Bruker, Massachusetts, USA). Images were acquired over 360° at rotation steps of 0.2 degrees. Exposure was set to 590 ms per frame, and source voltage and current values were 49 kV and 200 μA, respectively. The data were reconstructed with Nrecon software (version 1.6.6.0, SkyScan). Reconstruction duration was 1.48 s per slice, with ring artifact correction set to 6. Image pixel size was 8.3 μm.

## Results

### Evaluation of fluid velocity measurements in a flow phantom

The first experiment aimed to assess the sensitivity of the convection-MRI sequence to low-velocity flow, using a simple flow phantom. Velocity measurements, with the vascular nulling preparation switched off and using a *v_enc_* of 1000, 2000 and 5000 μm s^−1^, in separate experiments. Theoretical velocity predictions were estimated from the known fluid inflow rate and phantom diameter, and assumed laminar flow profiles. Fluid flow was directed from the point of inflow of the phantom, to the point of outflow (left to right in the image in **Figure 1a**), and was greatest in a channel through the centre of the phantom (**Figure 1b**).

For *v_enc_* of 2000 and 5000, velocity measurements were significantly correlated with theoretical velocity predictions (p<0.01, Spearman’s rho), for a range of physiological inflow values (**Figure 1**). At lower *v_enc_* values, aliasing occurred at higher velocity values (resulting in significantly lower velocity estimates; denoted by arrows on **Figure 1c**), and signal crushing by velocity gradients (particularly at *v_enc_* ≤ 500 μm s^−1^). At the largest inflow rate (6.85 mL min^−1^, theoretical velocity 1000 μm s^−1^), significant spatial wrapping was found in raw velocity encoded data, for the majority of *v_enc_* values used, so this data point was disregarded.

Whilst, for *v_enc_* ≥ 2000 μm s^−1^, convection-MRI velocity measurements were significantly correlated with theoretical estimates, they displayed a slight overestimation (15 ± 23 %, based on the y-intercept of the regression line) (**Figure 1c**). The minimum flow rate that could be measured before the signal approached the noise background level, was 0.07 mL min^−1^ (corresponding to a velocity of 20 (μ s^−1^).

### Evaluation of vascular nulling in tumors

The convection-MRI sequence was next evaluated *in vivo*, using subcutaneous mouse tumor xenograft models (SW1222 and LS174T human colorectal carcinoma cell lines). Our first objective was to assess the efficacy of the vascular nulling component. As the preparation aims to destroy the signal from fluid flowing in blood vessels, taking the ratio of images acquired with and without vascular nulling (the nulling ratio) allowed us to approximately estimate the vascular blood volume (23). In subcutaneous colorectal xenograft models (SW1222 and LS174T, n = 4 for each), the measured nulling ratio ranged from 0 to 80% and was heterogeneously distributed throughout the tumors (**Figure 2a**). On average, vascular nulling induced a decrease in signal of 18% in SW1222 and 13% in LS174T tumors, a difference that was significantly different (*p* < 0.05, Mann-Whitney). No nulling was observed in a static agar phantom, as expected, due to the absence of flowing fluid (**Figure 2a**).

**Figure 2.**
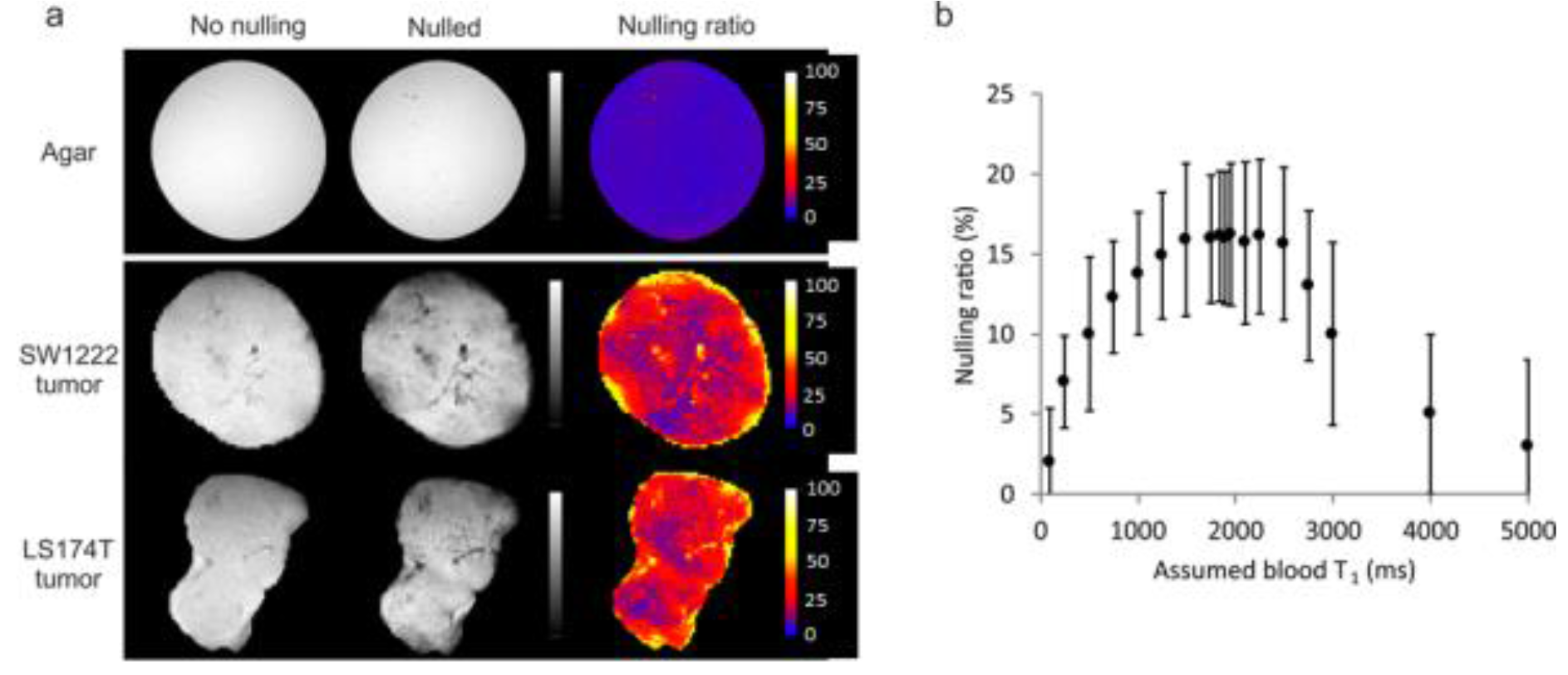
Evaluation of vascular nulling in tumor xenograft models. (a) Example maps showing the nulling ratio (the ratio of images acquired with vascular nulling to one acquired without vascular nulling) in an agar phantom and two different tumors. In the agar phantom, the nulling ratio was zero (top row), as expected due to the absence of flowing fluid. (b) A plot of the average nulling ratio as a function of the assumed blood longitudinal relaxation time (*T*_1,blood_). The assumed value of *T*_1,blood_ is used to set the recovery time following the inversion preparation (*t*_rec_ = ln(2) *T*_1,blood_). The graph shows that, for a range of *T*_1,blood_ of 1600 to 2500 ms, the nulling ratio is maximal. At lower values, the nulling is lower as the signal from blood has not recovered to a null point; at larger values, the signal has recovered past the null. The plateau represents a region where the signal from blood is near to or at the null point and has sufficient time to flow into and replace unlabelled blood within the imaging slice.

Plotting the mean nulling ratio as a function of the assumed blood longitudinal relaxation time (*T*_1,blood_), from *in vivo* experiments in SW1222 tumors (n = 4), revealed a plateau centred at 1900 ms at which the signal decrease caused by nulling was maximal (**Figure 2b**), suggesting that vasculaing nulling was most effective for an assumed *T_1,blood_* = 1900 ms (and so was used throughout the remainder of the study).

### Measurement of fluid velocity in tumors

Complete sets of convection-MRI data (i.e. with both vascular nulling and velocity encoding) were next acquired in SW1222 and LS174T tumors (n = 6 for both) (**Figure 3**). Plotting fluid velocity vectors for each pixel in the tumors revealed macroscopic flow patterns (**Figure 3e** and **l**), which often extended across the entirety of the tumor. These patterns were further visualised using a streamlining algorithm (**Figs. 3f** and **m**) that connected paths of coherent interstitial fluid flow, and revealed flow from a single or multiple sources within each tumor, towards the periphery. This source was generally located either towards the centre of the tumor or at the interface with the abdominal wall. Streamlines were often observed to be directed radially from the source, towards the outermost edge of the tumor. We measured fluid speeds of 10 to 220 μm s^−1^ (95^th^ percentiles), with mean fluid speed of 110 ± 55 μm s^−1^ and 170 ± 80 μm s^−1^ (mean ± standard error) for SW1222 and LS174T tumors, respectively. These velocity measurements are significantly greater than those associated solely with interstitial flow (typically < 1 μm s^−1^ (18)). An assessment of measurement repeatability and reproducibility is provided in the supplemental materials.

**Figure 3.**
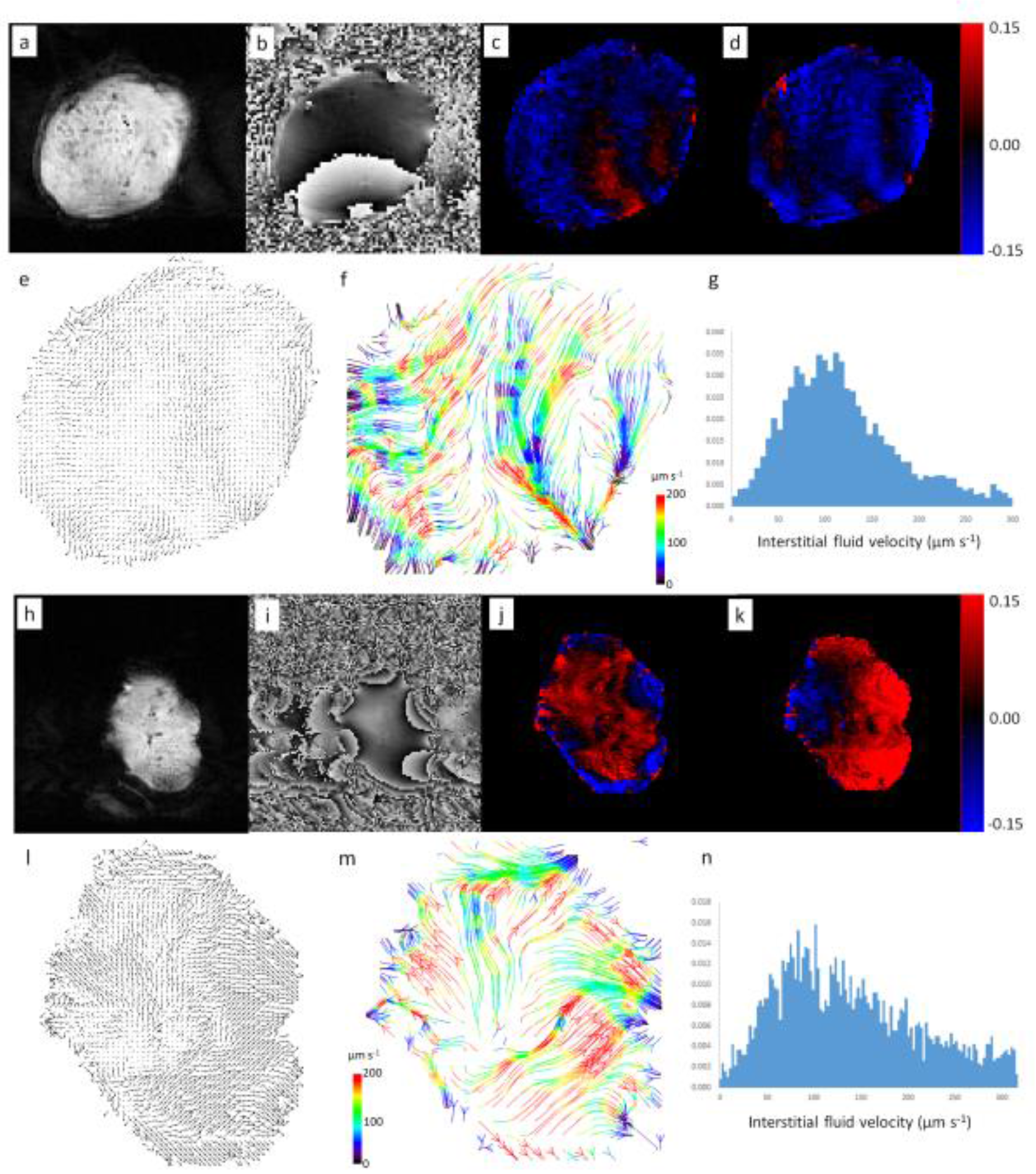
Example convection-MRI data sets (both raw image data and processed image data) in two example LS174T tumor xenografts: representative magnitude (a, h) and phase (b, i) images; maps of the change in phase with velocity-encoding gradients applied in vertical (readout) (c, j) and horizontal (phase-encoding) (d, k) directions. Velocity vector maps (e, l) show the direction of fluid transport through the tumor interstitium, which is better visualised using a streamlining algorithm (f, m) to connect pathways of coherent fluid convection (bottom row, colored arrows). Streamlines are color-coded to reflect the local fluid speed, which is also represented in histograms (g, n).

### Numerical simulations of tumor fluid dynamics

Monte Carlo simulations of tumor fluid dynamics were performed using physiological measurements previously reported in the literature. **Figure 4a** shows a schematic diagram of the simulations. The Hagen-Poiseuille equation used in these simulations estimated fluid velocities in blood vessels in the range of 0.2 to 15 mm s^−1^, which contrasts with the much slower fluid velocities in the interstitium (explicitly varied between 1 and 100 μm s^−1^).

**Figure 4.**
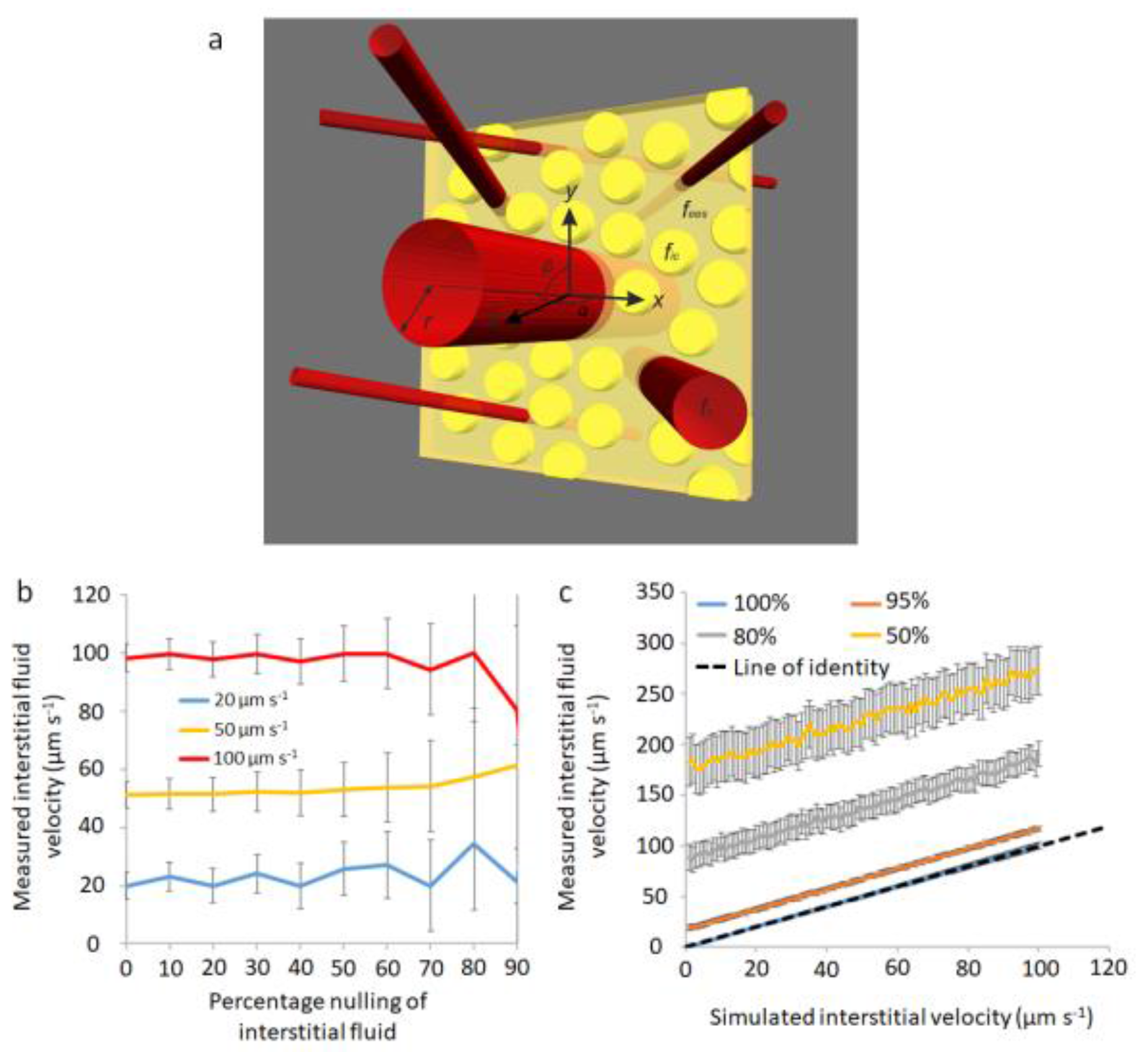
Results of simulations of fluid flow in SW1222 tumors. (a) shows a schematic diagram of the multi-compartment simulations (not to scale), in which red tubes represent blood vessels, yellow spheres are cells and the yellow cuboid represents the imaging slice. Parameters from the numerical simulation are overlaid. In (b), the mean value of IFV is plotted as a function of the percentage nulling of the interstitial fluid, for three simulated velocity values (0.02, 0.05 and 0.1 mm s^−1^). Error bars show the standard error in the mean value from 1000 Monte Carlo simulations. The graph in (c) shows IFV plotted against simulated interstitial velocity, for four vascular fractional nulling values (100, 95, 80 and 50%). Error bars show the standard error in the mean.

In the first set of simulations, it was found that *n_v_* was consistently greater than 99.9%, for each set of parameter values, suggesting that vascular nulling in this type of tumor is feasible. A fraction of the interstitial fluid was explicitly nulled to represent leakage of nulled fluid from the vasculature, and unintentional nulling of the interstitial fluid signal. Under these conditions, the accuracy of IFV measurements was unaffected by the nulling of interstitial fluid (Figure 4b), but precision progressively decreased with increased nulling (evidenced by the increasing standard error shown in **Figure 4b**). This loss of precision was caused by the decrease in signal-to-noise ratio resulting from the nulling, although the value of the simulated IFV had no influence on the magnitude of this effect.

In the second set of simulations, a progressive, incomplete nulling was imposed on the vascular fluid. It was found that both the accuracy and precision of IFV measurements became poorer as the fractional nulling of vascular fluid decreased (**Figure 4c**). The mean relative error in velocity measurements was 15%, 50% and 115% for 95%, 80% and 50% vascular nulling, respectively. As shown in **Figure 4c**, incomplete vascular nulling introduced a progressive offset to velocity measurements, but the gradient of each line is constant.

### Estimation of fluid pressure and comparison with a direct measurement with a pressure transducer

Convection-MRI provides measurements of fluid velocity vectors (i.e. with both direction and magnitude), from which we inferred an effective pressure (*P*_eff_) using Darcy’s Law of fluid flow in a permeable medium. Mean *P*_eff_ from convection-MRI measurements was 13 ± 2 mm Hg in SW1222 tumors and 16 ± 5 mm Hg in LS174T tumors, which were not significantly different (*p* > 0.05). Plotting the mean effective pressure value from the centre of each tumor, measured with convection-MRI, against those from pressure transducer measurements at corresponding locations, revealed a significant correlation (r^2^ = 0.77, *p* < 0.05, **Figure 5c**). On average, convection-MRI pressure measurements from the centre of tumors were 26% larger than those from the pressure transducer, and this over-estimation was larger at low IFP values. In these calculations, a fixed value was used for the hydrostatic conductivity (4.13×10^−8^ cm^2^ mm Hg^−1^ s^−1^ (4)).

**Figure 5.**
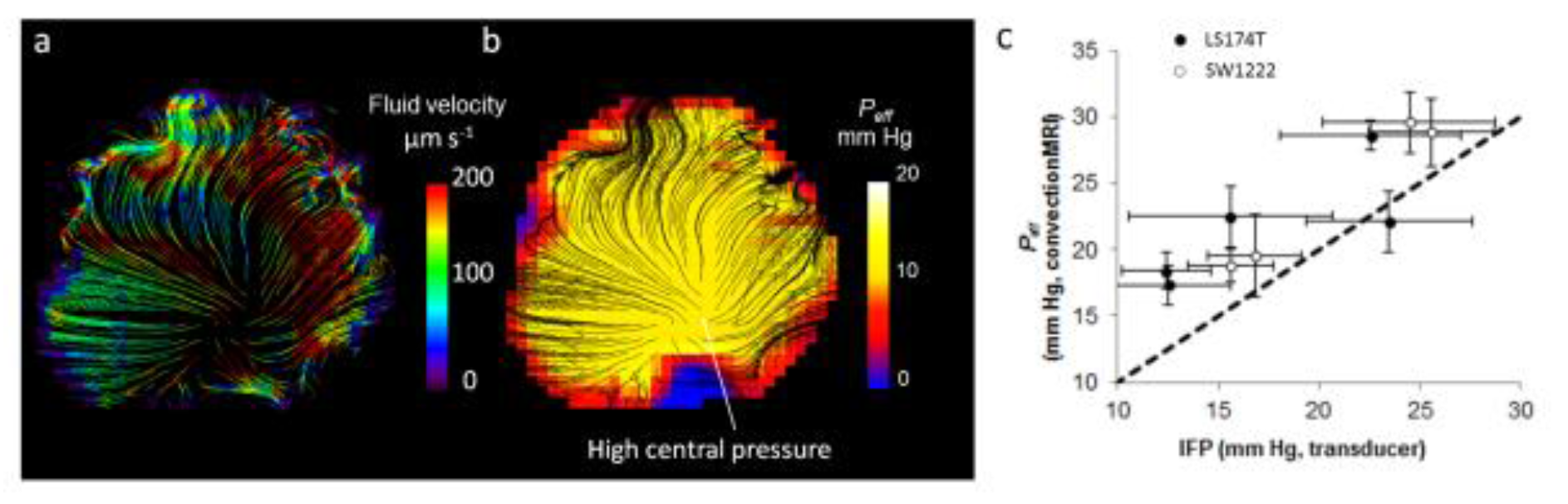
Estimation of effective fluid pressure (*P*_eff_) from convection-MRI measurements. a) An example convection-MRI fluid velocity streamline map and b) the corresponding effective pressure map from the same SW1222 tumor. c) Measurements of mean effective fluid pressure with convection-MRI, *vs* direct measurement of interstitial fluid pressure with a pressure transducer. Each point corresponds to the mean pressure in a different tumor, and error bars represent the standard error. Convection-MRI and pressure transducer measurements were significantly correlated (*p* < 0.05, Spearman’s rho).

### Comparison of tumor blood flow and fluid velocity distributions

Next, the coupling of intravascular and interstitial flow was probed, in a set of experiments in which measurements of tumor fluid velocity were acquired with convection-MRI, alongside measurements of vascular perfusion from arterial spin labelling (ASL) MRI. These were accompanied by high resolution images blood vessels acquired *ex vivo* from vascular casts at high spatial resolution (6 μm), with micro-CT.

Our first observation was that regions of high vascular perfusion (**Figure 6a**) measured *in vivo* coincided with highly vascular regions at the periphery of both tumor types, as assessed by microvascular casting (**Figure 6b**, **Supplemental Movie 2** (https://goo.gl/o9dE6g), **Supplemental Movie 3** (https://goo.gl/Uss2iz), **Supplemental Movie 4** (https://goo.gl/KoNc81)). Vascular perfusion was significantly greater in SW1222 tumors than in LS174T tumors (0.28 / 0.16 mL g^−1^ min^−1^ and 0.15 a 0.11 mL g^−1^ min^−1^, respectively, p < 0.01, Wilcoxon rank sum). This is consistent with the known vascular characteristics of each tumor type (29,30). Furthermore, as shown in the examples in **Figures 6b**, fluid velocity streamlines were found to emanate from highly perfused regions at the periphery of the tumors and follow paths through the interstitium that avoid other vascular regions. In the case of the example LS174T tumor shown in **Figure 6b**, interstitial fluid streamlines converged at the centre of the tumor and exited at a single, avascular point at the tumor periphery.

**Figure 6.**
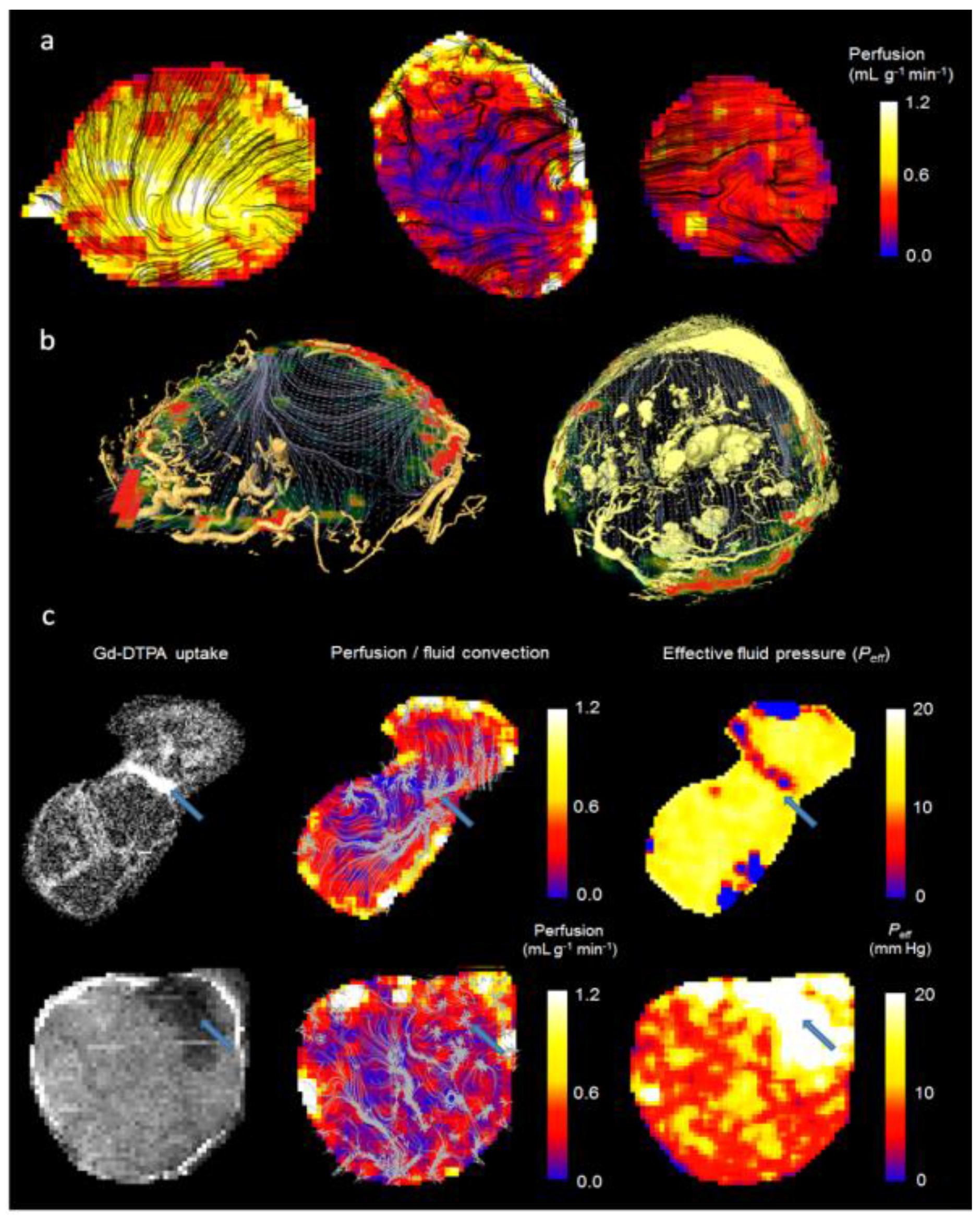
Comparison of perfusion (measured using arterial spin labelling) and convection-MRI measurements. (a) Perfusion maps in SW1222 tumors (left and right) and an LS174T tumor (centre). ASL measurements are shown as heatmaps, and are overlaid with black streamlines showing the measured path followed by fluid within tumors. (b) Fluid flow measured *in vivo* with convectionMRI are shown as grey streamlines and perfusion is shown as a colorscale. The location of blood vessels is represented by yellow volume renderings, acquired using *ex vivo* microvascular casting and imaged with micro-CT. Larger masses in the centre of the tumor correspond to swollen vessels; it is unclear if these were artificially distended during casting or if they were swollen under normal physiological conditions. Data are shown in example LS174T (left) and SW1222 (right) tumors. In the SW1222 tumor, larger vessel structures in the centre of the tumor could be due to swelling by the casting material, although it is unclear if these vessels would have also been swollen under normal physiological conditions. (c) The relationship between fluid flow, vascular perfusion and the delivery of a medium molecular weight contrast agent in two LS174T tumors. Left column: Uptake of Gd-DTPA, an MRI contrast agent; middle column: vascular perfusion maps (color scale) overlaid with interstitial convection streamlines (grey); right column: effective pressure measurements from fluid mechanical modelling of convection-MRI data. The blue arrow in the top row shows a region between two lobes of the tumor in which the contrast agent preferentially accumulates, interstitial convection streamlines converge, a limited vascular supply is evident, and has a low IFP. In the bottom row, a blue arrow highlights a region with limited contrast agent uptake, a limited vascular supply, and with raised IFP. Both examples show the ability of convection-MRI and ASL, in combination, to identify regions that preferentially accumulate or resist the accumulation of exogenously administered agents. This could potentially be extended to the prediction of the uptake of therapeutic agents.

The relationship between blood flow and interstitial convection was also explored by administering an exogenous contrast agent (gadolinium-DTPA) that could be independently visualised with MRI, and which is delivered via the tumor vasculature. The top row (first image) of **Figure 6c** shows an example LS174T tumor, in which raised accumulation of contrast agent can be observed between the two lobes that constituted the tumor (an animated version is provided in **Supplemental Movie 5** (https://goo.gl/PmfDCJ)). Streamlines from convection-MRI measurements revealed interstitial flow in opposing directions through the two lobes, which converged at their common boundary (**Figure 6c**, top row), corresponding to the region of raised contrast agent accumulation (blue arrow). According to a separate ASL measurement, vascular perfusion in this region was low, thereby implicating it as a sink for interstitial drainage, rather than the contrast agent being supplied directly by the tumor vasculature. In the bottom row of **Figure 6c**, data from a second LS174T tumor is shown, in which a region with poor contrast agent uptake is demarked by a blue arrow, with raised effective pressure. This therefore describes an opposite scenario to that of the first tumor, in which a region of raised IFP is potentially inhibiting the uptake of the contrast agent. Both of these examples demonstrate the potential of convection-MRI and ASL, in combination, to describe the uptake of exogenously administered agents (which could potentially be extended to therapeutic agents).

### The relationship between vascular perfusion and effective fluid pressure during tumor growth

The mechanisms that initiate and maintain raised IFP in tumors are of particular interest, and so we performed a further experiment that aimed to measure fluid velocity, effective fluid pressure and perfusion in tumors during growth from a small size (10 days following inoculation, defined as day 0), for a total of 10 days (**Figure 7a**). We found no significant difference in the mean growth rates for SW1222 and LS174T tumors, which were measured from tumor volume estimates using high-resolution T_2_-weighted MRI. Investigating the relationship between each parameter and tumor volume revealed a significant inverse correlation between vascular perfusion and tumor volume, whilst fluid velocity and effective pressure were both positively correlated with tumor volume (*p* < 0.05, **Figures 7b**). As necrosis is generally more evident in LS174T tumors than in SW1222 tumors, it might be expected that the measured apparent diffusion coefficient (ADC) would be correspondingly larger; however, there was no significant difference between the two tumor types, which is consistent with our previous measurements (30).

**Figure 7.**
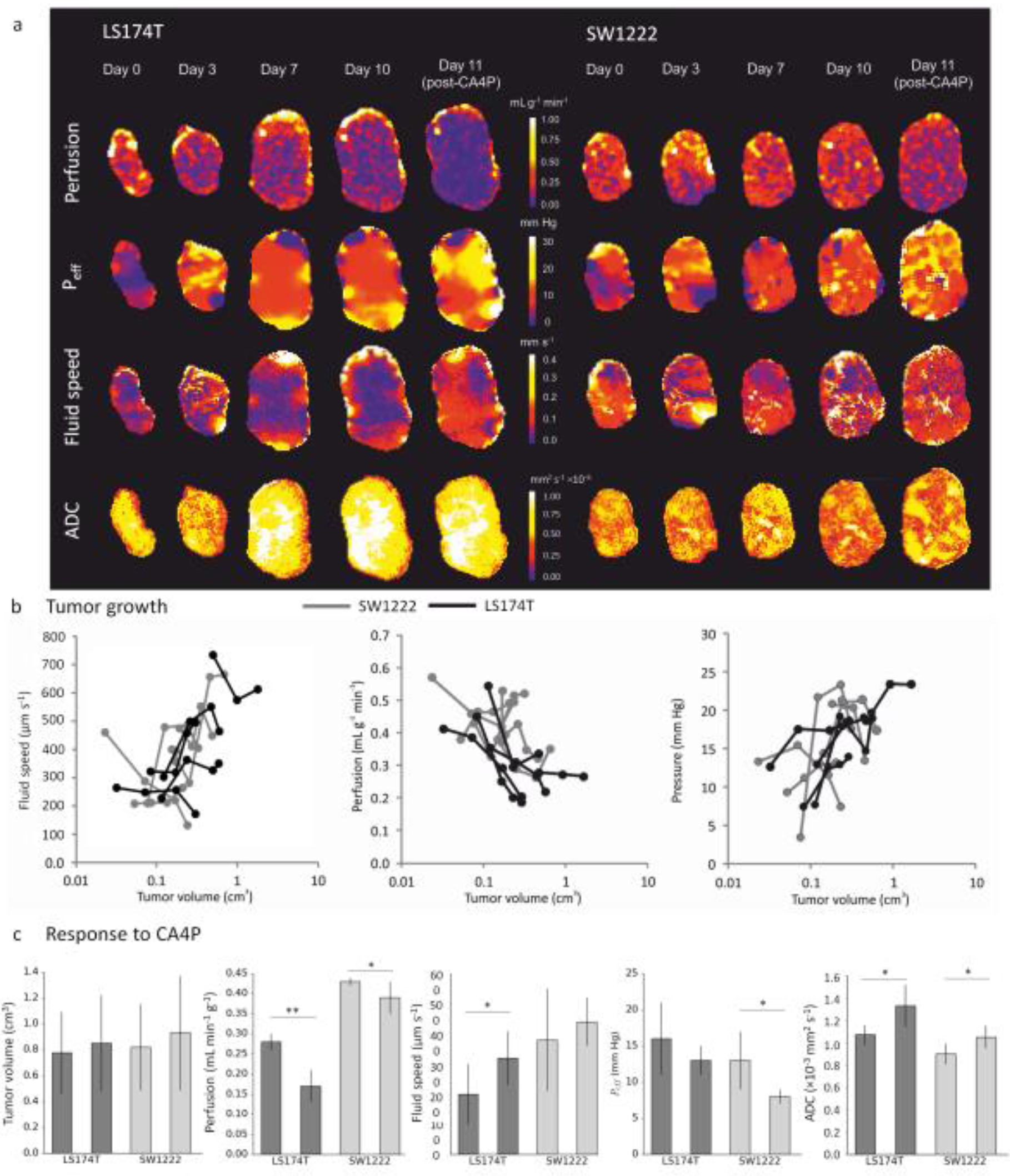
Tumor growth and response to therapy characterised with convection-MRI. (a) Example maps of vascular perfusion, effective pressure (*P*_eff_), fluid velocity and apparent diffusion coefficient (ADC) during 10 days of growth in two colorectal tumor xenografts (LS174T and SW1222), and at 24 hours following a single dose of the vascular disrupting agent CA4P (100 mg kg^−1^). (b) Scatter plots of the mean values of these parameters, from the whole tumor cohort. Fluid speed, measured using convection-MRI, tends to increase with tumor volume), whilst perfusion and *P*_*eff*_ decrease. Each point corresponds to a single measurement from a tumor, and black and grey lines connect individual tumors (LS174T (*n* = 6) and SW1222 (*n* = 7) colorectal tumor xenografts, respectively). (c) shows the mean change in each parameter at 24 hours following treatment with CA4P. Tumor volume did not significantly change with treatment, whilst in LS174T tumors, perfusion decreased and interstitial fluid speed and ADC increased. In SW1222 tumors, perfusion and *P*_*eff*_ significantly decreased and ADC significantly increased. Error bars represent the standard error in the mean; ** denotes p<0.01, * denotes p<0.05.

### Assessment of response to therapy with a vascular disrupting agent

At the end of the growth experiment, and immediately following the final MRI measurement, we administered a single dose of the vascular disrupting agent CA4P (100 mg kg^−1^) to all mice. Previous studies have shown that this rapidly induces vascular occlusion and haemorrhage (41), and a corresponding change in perfusion and IFP (42). With this in mind, we measured the change in perfusion, *P*_eff_ and fluid velocity at 24 hours following dosing. As expected, tumor volume did not change significantly with CA4P dosing, whilst perfusion significantly decreased (p<0.01 in LS174T and p<0.05 in SW1222). Fluid velocity significantly increased in LS174T tumours (*p<*0.05)) and *P*_eff_ significantly decreased in SW1222 tumors (p < 0.05) (**Figure 7c**). Tumor ADC significantly increased at 24 hours after dosing, in both tumor types, presumably due to the onset of necrosis (*p* < 0.05, data not shown).

## Discussion

We have proposed here a new technique named convection-MRI, which aims to measure low-velocity fluid flow in tumors, using phase contrast MRI, by first nulling the fast-flowing intravascular signal with a dual-inversion preparation. The aim of this study was to evaluate whether convection-MRI could directly measure the velocity of interstitial fluid. In order to achieve this a number of assumptions associated with the measurement would need to be validated, and we aimed to evaluate the most significant of these in this study. We then used convection-MRI to study low-velocity fluid flow in tumor xenograft models, which appeared to reveal macroscopic flow patterns that were consistent with the flow of fluid through the tumor interstitium, albeit at velocities greater than those normally associated with interstitial flow.

In tumor xenograft models of human colorectal cancer, we calculated mean fluid velocity estimates of 110 ± 55 μm s^−1^ and 170 ± 80 μ s^−1^ in SW1222 and LS174T tumors, respectively, which is larger than those recorded in tumors using other techniques. A review by Munson *et al* reports IFV values of the order of 0.1 to 55 μm s^−1^, measured using a range of techniques, both in mouse tumor models and human patients (18). On this basis, the slowest flowing components of interstitial fluid would appear to be inaccessible with this technique, particularly as our flow phantom measurements could detect a minimum velocity of 20 μm s^−1^. It is worth also noting that measurements of the lowest velocity components (i.e. around 20 μm s^−1^) will be associated with a greater uncertainty, due to the influence of noise. The presence of fluid velocities greater than 55 μm s^−1^ in our *in vivo* measurements is potentially due to contamination by faster-flowing vascular fluid. This therefore presents a challenge for the interpretation of convection-MRI data, as it cannot reasonably be claimed that velocity measurements are entirely interstitial in origin, nor that the entire range of interstitial flow velocities can be captured.

These observations were further explored through Monte Carlo simulations of a SW1222 tumors, which predicted vascular nulling of greater than 99.9%, for all sets of physiological parameters considered. They also showed that un-nulled vascular fluid can, in principle, be entirely replaced by nulled fluid within the recovery time *t_rec_* between vascular nulling and velocity encoding. In the simulations, straight tumor vessels were modelled, whilst real tumor vessels are likely to take more tortuous paths, thereby potentially explaining the disparity between our simulation results and the fluid velocities measured *in vivo* (43). Sensitivity to these slower flowing components (and improved specificity towards interstitial flow) could therefore potentially be achieved by improving vascular nulling for slow-flowing blood, and increasing the acquisition *v_enc_*.

Whilst not an explicit component of the convection-MRI acquisition, *in vivo* estimates of blood volume were also derived by comparing nulled and un-nulled images. This analysis provided blood volume estimates that were greater than, but of the order of the microvascular density previously reported for these tumor types. Previous studies provide average blood volumes of between 9 (44) and 25% (30) for SW1222 and 8 (30) to 9% (45) for LS174T tumors, whereas we measured an average percentage vascular nulling of 18% and 13% for SW1222 and LS174T tumors, respectively. The largest blood volume values were measured at the tumor periphery (of the order of 80%), where vessels tend to be larger and more concentrated (29), which demonstrates the ability of convection-MRI to capture the spatial heterogeneity of tumor vasculature. These data could therefore act as a useful complement to the convection-MRI acquisition, although requires the acquisition of the additional un-nulled image.

Despite our velocity measurements being greater than those previously reported in the literature, we still found a significant correlation between *P*_eff_ (derived from the spatial derivative of velocity measurements) and measurements of IFP measured directly with a pressure transducer, although with a 26% overestimation (**Figure 5c**). We used Darcy’s Law to quantify pressure from velocity vector fields, which is widely used in the modelling of fluid dynamics in biological tissue. It is valid for systems with a low Reynolds number which, in tumors, is typically much less than 1 (46). When estimating absolute pressure from velocity vector fields, pressure boundary conditions are required, and here we assumed that the mean interstitial pressure at the periphery of tumors was maintained at atmospheric pressure (34). Equally, we used a fixed value for the hydrostatic conductivity of tumor tissue, *K*. *K* provides a linear scaling to pressure estimates (as per Darcy’s Law in equation 2), and so errors associated with this value will therefore introduce a linear offset into pressure estimates. As *K* can only currently be measured by invasive means, use of a fixed value is presently the only available solution. Our assessment of repeatability and reproducibility (36% and 59%) were slightly larger, but of the order of those measured for other MRI techniques (e.g. 13% - 62% for diffusion MRI (47) and 8.9 to 18.2 % for dynamic contrast-enhanced MRI (48)). This variation could potentially be reflective of true physiological changes over short timeframes.

We next probed the relationship between microvascular perfusion and interstitial fluid pressure, which has not been extensively studied, mainly due to a lack of available, non-invasive techniques. Our measurements, acquired over a period of ten days of tumor growth, revealed differences in the microenvironments of two colorectal tumor types. We found that perfusion on average decreased with tumor volume, whilst effective fluid pressure increased, which is in agreement with previous studies (20,49). Both low vascular perfusion and raised IFP can act as a barrier to drug delivery, our combined approach would allow these properties to be probed simultaneously and monitored longitudinally. Generally, LS174T tumors are more necrotic than SW1222 tumors, at 59 ± 4 % and 10 ± 3 %, respectively, in fully-grown tumors (50), and which likely increases to these levels throughout tumor growth. Large regions of central necrosis occurs at 24 hours following dosing with CA4P (51). However, the presence of necrosis should not affect the convection-MRI acquisition (instead, it could be of interest to study further, to investigate the influence of necrosis on interstitial fluid pressure and drug delivery).

We have also shown here that ASL and convection-MRI, in combination, can be used to assess response to the vascular disrupting agent CA4P. The decrease in *P*_eff_ that we observed in SW1222, and no significant change in LS174T tumours, is in agreement with the decrease or stasis found in other studies. It is of interest that SW1222 exhibited a minimal decrease in perfusion with CA4P treatment, compared with the response in LS174T, yet still exhibited a decrease in *P*_eff_ and increase in ADC. This could be due to complex changes in tumor fluid dynamics, which could be of interest to further explore.

A number of studies have compared other imaging-based parameters with pressure transducer measurements of IFP, such as dynamic contrast enhanced MRI (8,34) and/or diffusion MRI (52). These have generally shown a correlation with IFP, but such relationships are not necessarily causal and could rely on associations with other aspects of tumor pathophysiology. However, the uptake of contrast agent has previously been shown to correlate with the uptake of chemotherapeutic drugs (53) and to probe interstitial transport and lymphatic drainage (54), and convection-MRI could compliment these approaches via its ability to directly measure the flow of interstitial fluid within tumors. Some attempts have been made to quantify IFP from contrast agent studies (55,56), and a study comparing this approach and the convection-MRI technique would be useful. It is worth noting that our calculation of the nulling time within the convection-MRI signal model, assumed complete recovery of the longitudinal magnetisation within a practical acquisition recovery time. However, a more accurate estimate could be used that takes into account for incomplete recovery of the longitudinal magnetisation. Simulations using the Bloch equations suggest that this effect introduces small (approximately 3%) error into the vascular nulling efficiency (data not shown), but which could be utilised in future studies.

Translation of the convection-MRI sequence onto clinical systems should be straightforward, although requires further investigation and numerical simulations. Factors such as signal-to-noise and resolution differences between the high-field experimental system used here and that of clinical MRI systems would need to be explored. Equally, whether the approach offers sufficient sensitivity to the slowest fluid flows in the clinical setting would need to be evaluated, and tumor motion during scanning would need to be minimised. Conversely, *T*_1_ is shorter at clinical field strengths, allowing a shorter in-flow time during vascular nulling (although human blood *T*_1_ is longer than in mice).

## Acknowledgements

We acknowledge the support received for the Kings College London & UCL CR-UK and EPSRC Comprehensive Cancer Imaging Centre, in association with the MRC and Department of Health (England), (C1519/A10331). BBSRC/AstraZeneca Industrial Partnership Studentship (BB/E528979/1), the UK Regenerative Medicine Platform Safety Hub (MRC: MR/K026739/1), Eli Lilly and Company, the EPSRC (EP/N034864/1) and the Wellcome Trust (WT100247MA, 091763/Z/10/Z). We thank OXiGENE for supplying CA4P.

